# Epigenetic de-repression of basal cell metaplasia in aging AT2 cells is a risk factor for idiopathic pulmonary fibrosis (IPF)

**DOI:** 10.64898/2026.06.09.731212

**Authors:** Stefano A. Iantorno, Ying Wei, Kiana Garakani, Alexis N. Brumwell, Tsung Che Ho, Marylene Toigo, Jaymin J. Kathiriya, Asres Mitke, Vivianne Ding, Zea Borok, Johannes Kratz, Michael A. Matthay, Paul J. Wolters, Harold A. Chapman, Claude Jourdan Le Saux

**Affiliations:** Department of Medicine, Division of Pulmonary, Critical Care, Allergy, and Sleep Medicine, University of California San Francisco, San Francisco, California; Department of Stem Cell Biology and Regenerative Medicine, Icahn School of Medicine at Mount Sinai, New York, New York; Department of Medicine, Division of Pulmonary, Critical Care, Sleep Medicine, and Physiology, University of California San Diego, San Diego, California; Department of Surgery, Division of Thoracic Surgery, University of California San Francisco, San Francisco, California

## Abstract

Idiopathic pulmonary fibrosis (IPF) is a fatal, age-associated lung disease in which alveolar type II (AT2) cells lose regenerative capacity and can adopt aberrant basal-like fates that promote fibrosis. Using 3D organoid co-cultures with primary human fibroblasts, we find that healthy human AT2 cell trans-differentiation into KRT5+/KRT17+ basal cells increases progressively with age, while differentiation into RAGE+ AT1-like cells decreases. We identify a shared gene signature in AT2 cells at downstream targets of p63 characterized both by acquisition of bivalent, poised chromatin marks with age and increased accessibility in IPF, indicating epigenetic “priming” towards a basal cell lineage. *In vitro* treatment of young AT2 cells with IL-1β recapitulates this priming toward basal differentiation via a NF-kB-regulated histone demethylase, JMJD3. Conversion of primed AT2 cells to a basal fate requires recruitment of a shared transcription factor, KLF5, from AT1-specific to basal-specific promoters by HIF-1α. AT2 cells instead convert to KRT5-/KRT17+ basaloid cells via a non-age-dependent pathway that requires KLF5-SMAD2/3 complexing through TGFβ1 signaling. These findings define an inflammation-driven epigenetic de-repressive mechanism that links aging, inflammatory stress, hypoxia, and dysfunctional epithelial metaplasia, and accounts for the likely origin of aberrant epithelial cell populations in fibrotic lung disease.

## Introduction

A growing aging population will have a significant impact on health care systems worldwide in the coming decades^1^, underscoring the importance of developing new scientific insights and management strategies for age-related chronic diseases such as idiopathic pulmonary fibrosis (IPF). Advanced age is the most important risk factor for IPF, which has an average age of onset of 63 years^2,3^. Over 40,000 new cases of IPF are recorded yearly in the US, with an expected survival of 3-5 years from the time of diagnosis^4,5^. Few treatment options are available for eligible patients, and include antifibrotic therapies, which can slow lung function decline, and lung transplantation^6^. However, both are associated with significant side effects and adverse events, and do not represent curative options, highlighting the need for better management strategies.

Genetic studies of familial forms of IPF point to causative mutations affecting alveolar type II (AT2) cell function, such as surfactant production, as key initiators of disease^7,8^. AT2 cells act as dedicated alveolar epithelial stem cells that both self-renew and differentiate into alveolar type I cells (AT1), which instead represent the main structural components of the alveolus and interface with capillaries to allow diffusion of oxygen and carbon dioxide. A long-standing paradigm of IPF pathogenesis suggests that AT2 cell senescence and/or apoptosis, likely due to impaired cellular responses to chronic stress or injury^9–11^, predispose to fibrosis by reducing the regenerative potential of this crucial stem cell niche, thus disrupting the ability of the alveolar epithelium to regenerate itself and facilitate gas exchange. The cytopathologic correlates of this process are found in the appearance of a Usual Interstitial Pneumonia (UIP) pattern in histology section from the distal lung^12,13^. This pattern is characterized by the presence of subpleural microcysts and “honeycomb” cysts lined with aberrant epithelial cells, including ectopic secretory and basal cells, typically seen overlaying fibroblastic foci and juxtaposed to areas of normal-appearing alveoli. The extent of UIP histopathologic pattern correlates with disease severity and outcomes^14^, pointing to ongoing replacement of AT2 and AT1 cells by airway-like aberrant epithelial cell populations as an important feature of disease progression^15–18^.

Expansion of ectopic airway-like cell populations in human disease is thought to be driven by mobilization of distal airway basal cells, as suggested by murine models of lung injury and fibrosis^19–24^. Recent studies from our group and others^25–28^, however, have shown that primary human AT2 cells can trans-differentiate in response to fibroblast-derived factors into KRT5+/KRT17+ basal and KRT5-/KRT17+ basaloid cells that resemble the aberrant epithelial cell populations that have been described in several forms of fibrotic lung disease, including IPF, with single-cell RNA-seq approaches^29,30^. Several of these studies have highlighted the importance of hypoxia, or HIF signaling, in driving this metaplastic process^31–33^. Notably, AT2 cells in murine models of lung disease display significant less plasticity than primary human AT2 cells^34^, pointing to important differences in the underlying biology. We therefore sought to account for the increased risk for IPF seen with age by investigating the possibility that metaplastic trans-differentiation of primary human AT2 cells into basal-like cells may also be a function of aging. We hereby characterize an aging-dependent, inflammatory-mediated epigenetic mechanism in AT2 cells that leads to de-repression of a gene signature marked by both activating H3K4me3 and repressive H3K27me3 histone marks at their promoter sites, a chromatin state that has been described as bivalent, or poised^35–38^. This poised state predisposes aged AT2 cells to acquire basal cell features via the HIF1α-dependent recruitment of KLF5 transcription factor complexes from AT1-specific to basal-specific binding sites, thus suppressing AT1 regeneration. This mechanism represents a new model for how cellular aging in epithelial stem cells may directly contribute to disease, in this case fibrogenesis, by promoting the appearance of dysfunctional cell fates in response to differentiation signals via prior epigenetic “erosion”^39,40^ of their pre-programmed lineage commitment.

## Results

### AT2 metaplasia during *in vitro* trans-differentiation in 3D organoid co-culture with primary human mesenchyme is positively correlated with donor age

We generated standard 3D alveolar organoids as previously described, using freshly isolated AT2 cells randomly paired with lung fibroblasts from each newly available non-diseased donor lung tissue obtained through UCSF. Colonies were harvested after 14 days of AT2 cell-mesenchymal co-culture, fixed and embedded, and stored frozen without evaluation. After 18 months, the accumulated OCT-embedded organoids were stained by immunofluorescence for AT2 (SFTPC), AT1 (RAGE), and basal cell (KRT5) markers, and evaluated as a single batch by two observers who were blinded to the age of each sample (Figure 1A). Both AT2 and fibroblast donors in this cohort included both sexes. Additional organoids were similarly generated from donors and the EPCAM+ epithelial cell fraction was processed for RNA extraction at the same time point. Surprisingly, the fraction of colonies containing at least 25% KRT5+ epithelial cells (hereafter termed “KRT5+ colonies”) increased nearly linearly with AT2 cell donor age (Figure 1B, 1C). By donor age 70, the fraction of KRT5+ colonies was close to 100% in most of the donors. Conversely, the fraction of colonies marked by at least 25% expression of RAGE (“RAGE+ colonies”), a canonical marker of AT1 cells that corresponds to the *AGER* gene, showed a striking inverse correlation with AT2 cell donor age (Figure 1B and D). Notably, colonies generated from AT2 cell donors at the older range of the spectrum did not have appreciable staining for either RAGE or for SFTPC after 14-days of fibroblast co-culture, while there was significant positivity for both markers in colonies from younger AT2 cell donors. We then examined expression of basal cell markers (*KRT5*, *KRT17*, *KRT19*, *TP63*) and AT1 markers (*AGER*, *CLIC5*, *PDPN*) by qPCR in EPCAM+ epithelial cells at 14 days of organoid co-culture. Donors were grouped into “Young” or “Old” categories by using a cutoff of donor age <51 and ≥51, respectively. Interestingly, *KRT17* mRNA expression was not significantly different between “Young” and “Old” categories, while *KRT5*, *TP63*, and *KRT19* mRNA levels were significantly increased in the “Old” category (Figure 1E, Supplementary Figure 1); *AGER* and *CLIC5* conversely were significantly decreased in the “Old” category, while *PDPN* was not significantly different (Figure 1F, Supplementary Figure 1). We then considered whether the age of primary human fibroblast donors was a variable affecting AT2 cell metaplasia in this system. To test this, we measured correlation between the proportion of KRT5+ and RAGE+ colonies in each sample and the age of the fibroblast donor used for that sample. We found no significant correlation for either outcome (Figure 1G and H), suggesting that this differential propensity toward AT1 or basal cell fate is due to AT2 cell intrinsic factors. Collectively, these data indicate increased basal cell metaplastic potential and reduced AT1 cell trans-differentiation with advancing AT2 cell donor age.

**Figure 1.**
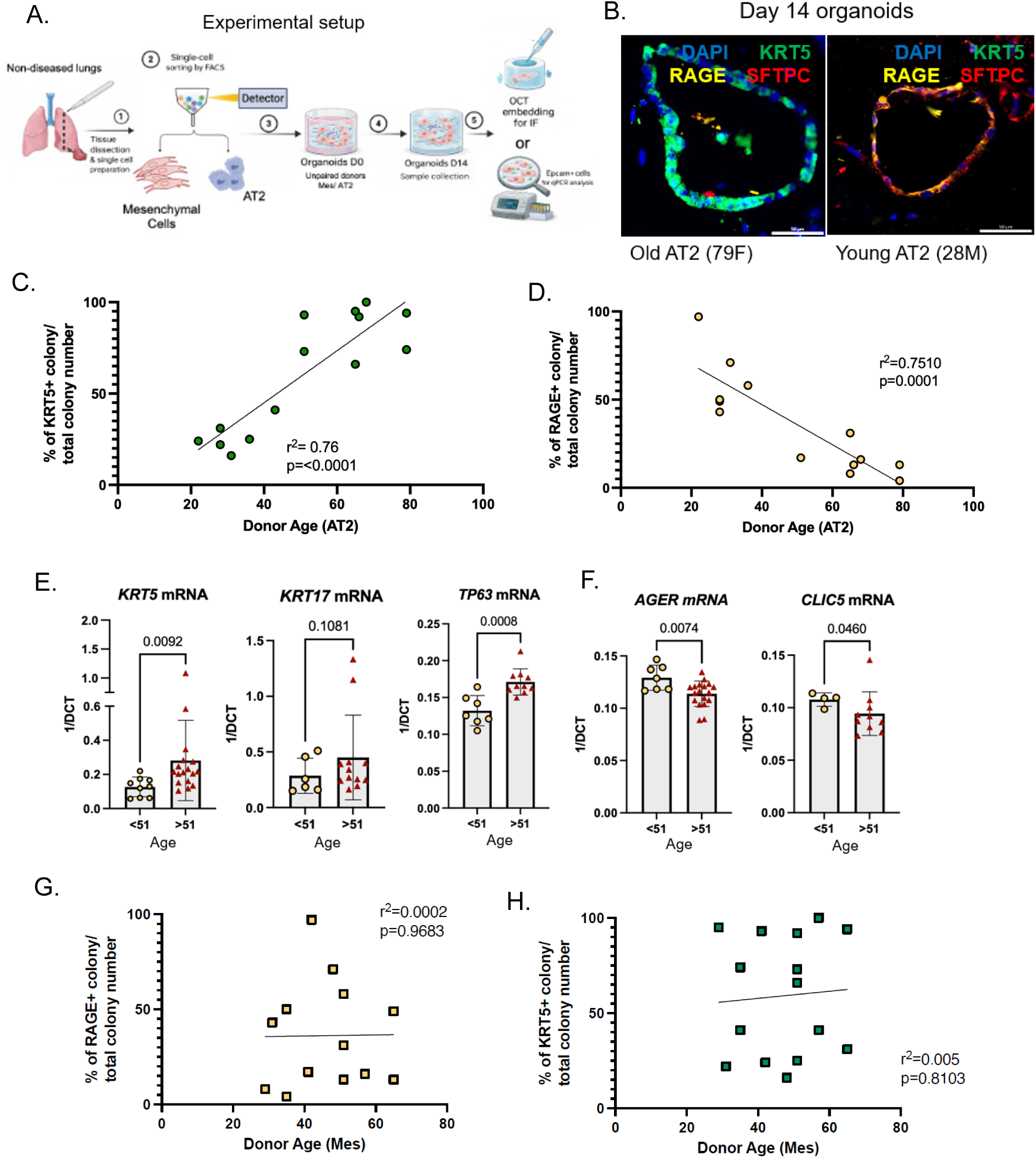
Primary human AT2 cells have increased propensity for trans-differentiation into basal cells with age. **(A)** Schematic showing experimental workflow. Briefly, AT2 cells were isolated from non-diseased donors with flow cytometry and organoid cultures were prepared with primary human fibroblasts. After fixing, OCT blocks were cut, stained, and analyzed by immunofluorescence microscopy as a batch by two blinded observers. Additional organoid cultures were used for qPCR analysis. **(B)** Representative images of day 14 organoid from “Young” (age <51) and “Old” (age >51) donors showing immunostaining for canonical basal (KRT5), AT2 (SFTPC), and AT1 markers (RAGE). **(C-D)** Proportion of KRT5+ vs RAGE+ colonies in day 14 organoids as a function of AT2 donor age using a 25% cutoff for positivity. Each data point represents a separate donor lung. There is a significant positive linear correlation with age for proportion of KRT5+ colonies, and a significant inverse correlation for proportion of RAGE+ colonies with age, as shown by R-squared value > 0.75 **(E)** Gene expression by qPCR for basal cell associated markers in day 14 organoids, split by age, showing significant increase in *KRT5* and *TP63*, but not *KRT17*, mRNA. **(F)** Gene expression by qPCR for AT1 associated markers (*AGER*, *CLIC5*) in day 14 organoids, split by age, showing significant decrease in mRNA levels with age of AT2 cell donor. **(G-H)** Proportion of KRT5+ vs RAGE+ colonies in day 14 organoids as a function of primary fibroblast donor age, showing no significant correlation. Statistical significance was determined by simple linear regression **(C, D, G, H)** and unpaired t-test **(E, F)**.

### A set of p63-associated, IPF-relevant genes acquire bivalent histone marks at their promoter sites with increased age in AT2 cells, suggesting an epigenetic poised state

To further elucidate mechanisms that might be controlling this age-dependent differential propensity in trans-differentiation capacity towards either a basal or AT1 cell fate, we performed bulk CUT&Tag sequencing to identify the location of two histone modification marks, histone H3 lysine 4 trimethylation (H3K4me3) and histone H3 lysine 27 trimethylation (H3K27me3) in primary human AT2 cells isolated from either “Young” or “Old” donors (n=3 for each age category, see Supplementary Table 1). Importantly, these marks are associated with either activated (H3K4me3) or repressed (H3K27me3) transcription. A chromatin state where these two marks co-exist, termed bivalent chromatin, has been described, and its role in maintaining genes in a poised state in multipotent stem cells until the appropriate developmental trigger occurs is well established in the literature^41–46^. We annotated genome-wide promoter regions, defined as a 6kb window around each transcription start site (TSS +/− 3000bp), marked by H3K4me3 alone (“activated”), H3K27me3 alone (“repressed”), and both H3K4me3/H3K27me3 (“bivalent”) by identifying consensus peak calls shared by all replicates in each condition (Supplementary Figure 2). Interestingly, *KRT5* and *KRT17* were among the set of genes marked by bivalent histone modifications at their promoters in AT2 cells from “Old” donors, while they only harbored repressive marks in these same genomic regions in “Young” donors (Figure 2A). We defined this transition from an epigenetic repressed to a poised state as age-dependent “de-repression”. Consistent with age-dependent de-repression, we observed an expansion of genes marked by bivalent promoters with increased age in AT2 cells, from 29% to 46% of total annotated genes, while the proportion of active and repressed genes decreased (Figure 2B). H3K4me3 enrichment around the transcription start site of these genes increased with age, while H3K27me3 decreased with age, pointing to demethylation of H3K27 and methylation of H3K4 residues at these loci (Figure 2C). Of note, the *AGER* locus did not appear to have consistent presence of either histone mark at its promoter across conditions, suggesting that epigenetic regulation by these two histone marks is not important for its expression, which would be compatible with a permissive or de-regulated epigenetic state.

**Figure 2.**
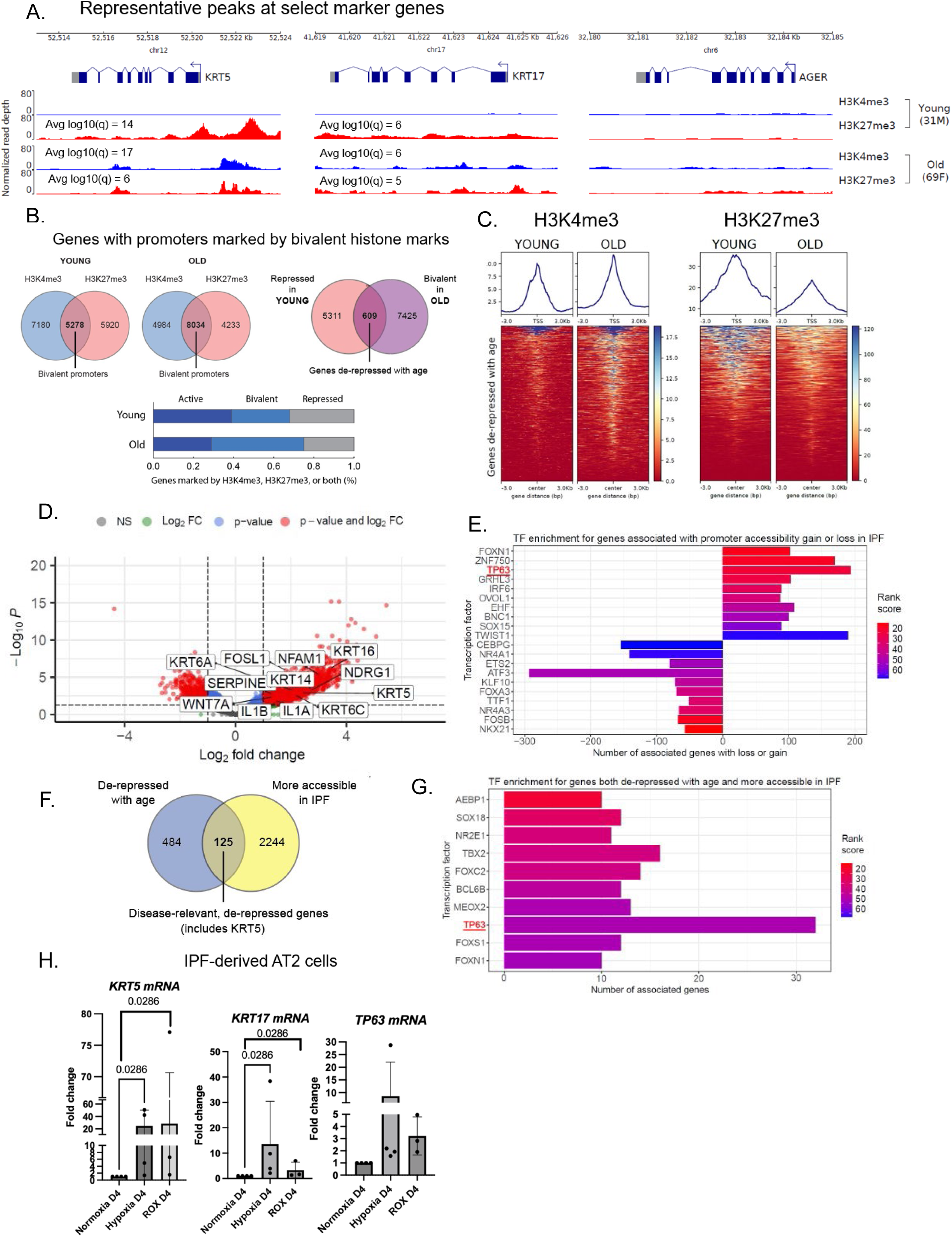
AT2 cells acquire a shared p63-enriched gene signature that is de-repressed with age and has increased accessibility in IPF. **(A)** Representative genomic tracks of normalized read depth from CUT&Tag sequencing for H3K4me3 and H3K27me3 histone marks in AT2 cells from donors of different ages at select marker loci. Both *KRT5* and to some extent *KRT17* promoters become bivalent, i.e. marked by both H3K4me3 and H3K27me3, with age, while they are repressed, i.e. marked by H3K27me3 only, in younger donors. Adjusted p-values (as average −log10(p)) are shown for any peak calls. *AGER* promoter did not have any peak calls. **(B)** Identification of genome-wide bivalent promoter sites (defined as TSS +/− 3kb) in donors of different ages. There is an expansion of bivalent promoters with age from 29% to 46% of total genes marked by histone modifications. **(C)** Enrichment around transcription start site (TSS) for H3K4me3 and H3K27me3 in genes that become de-repressed with age. There is significant decrease in enrichment for H3K27me3, and slight increase for H3K4me3. **(D)** Differential promoter accessibility in AT2 cells derived from IPF vs age-matched non-diseased donors. Highlighted are several relevant p63 target genes, all with significantly increased accessibility in their promoter region in IPF. **(E)** Gene set enrichment analysis using the ChEA3 transcription factor database for genes that have significant gain or loss in promoter accessibility in IPF-derived AT2 cells, as determined by the previous differential accessibility analysis. Lower rank score is associated with more significant enrichment. The *TP63* gene, coding for p63, is among the top 3 transcription factors by rank score and associated with the largest number of genes with promoter accessibility gain in IPF, suggesting activation of this pathway. Conversely, NKX2-1 has the most enrichment by rank score among genes with promoter accessibility loss in IPF, suggesting repression of these downstream genes. **(F)** Identification of 125 genes that both acquire bivalent marks with age in AT2 cells and become activated (i.e. have increased promoter accessibility) in IPF-derived AT2 cells. This core set of genes includes *KRT5* but not *KRT17* and represents disease-relevant genes that become de-repressed with age. **(G)** Gene set enrichment analysis for disease-relevant genes that are de-repressed with age confirms enrichment for p63 genes (32 of 125 genes, or 26%, are p63 target genes). **(H)** The metaplastic potential of AT2 cells from IPF donors was tested under normoxia, hypoxia (2% O_2_), and with HIF agonist roxadustat (ROX) during 4 days of culture in AT2 medium. Hypoxia and HIF agonism both induce *KRT5* and *KRT17*, and to a smaller extent *TP63* mRNA. Statistical significance for peak calls in **(A)** was determined with MACS3 by comparing the CUT&Tag signal with background local lambda via a local Poisson test, and q-values estimated with Benjamini-Hochberg procedure. The average significance value −log10(q) for all peaks in the promoter region is shown, if any peaks were called. Statistical significance for **(H)** was determined by one-way ANOVA.

We next attempted to identify epigenetic signatures that may be responsible for AT2 metaplastic potential. Using all 609 genes de-repressed with age as input, we did not identify important regulators of epithelial cell fate by gene set enrichment analysis using the ChEA3 database, suggesting that many molecular pathways affected by the epigenetic changes that occur with cellular aging may not be relevant to AT2 metaplasia as it occurs in IPF (Supplementary Figure 2). We therefore turned to bulk ATAC-seq, which quantifies chromatin accessibility via a modified Tn5 transposase binding to open DNA regions^47^, to identify AT2-specific genes that are in an “activated” epigenetic state in IPF. We performed bulk ATAC-seq on AT2 cells isolated from IPF and age-matched non-diseased donors (n=3 for each condition) and again focused our analysis on peak calls falling within promoter regions (defined as TSS +/− 3000bp). Differential accessibility analysis found that several genes associated with hypoxia (*NDRG1*, *SERPINE1*), inflammation (*IL1A*, *IL1B*, *NFAM1*, *FOSL1*), and basal cell identity (*WNT7A*, *KRT6A/C*, *KRT16*, *KRT14*, and *KRT5*, but not *KRT17*) were significantly more accessible in IPF-derived AT2 cells (Figure 2D). Gene set enrichment analysis revealed that a p63 signature, a master regulator of basal cell identity^48–51^ encoded by the *TP63* gene, is among the top ranked transcription factors associated with genes marked by increased promoter accessibility in IPF (Figure 2E). Of note, NKX2-1, a transcription factor associated with specification of alveolar cell fate during development^52^, was the top hit for genes with reduced promoter accessibility, while Forkhead family members, such as FOXN1 and FOXA3, appeared to have opposite roles at genes either silenced or activated in IPF, as determined by their chromatin accessibility. Finally, to identify disease-relevant genes that are de-repressed with age, we prioritized genes whose promoter regions were both differentially accessible in our ATAC-seq analysis and transitioned from a repressed to a bivalent state in our CUT&Tag analysis. Again, we found significant enrichment for p63-associated genes, including the canonical basal cell marker *KRT5* (Figure 2F and G). In order to test the trans-differentiation potential of AT2 cells isolated from donors affected by IPF, we performed short term culture of these cells in mesenchyme-free 3D conditions. We found that AT2 cells derived from IPF donors similarly had increased expression of *KRT5* but not *KRT17* mRNA at day 4 of culture compared to AT2 cells derived from age-matched unaffected donors (Supplementary Figure 1). Of note, *TP63* mRNA, while expressed at low levels, had significantly higher levels in IPF-derived AT2 cells compared to non-diseased donors, consistent with our ATAC-seq results. Treatment with HIF agonist roxadustat or by placing cells under hypoxic conditions (2% oxygen) for 4 days led to strong induction of basal cell markers, confirming the poised status of these cells (Figure 2H). In summary, our integrated analysis points to priming of p63 binding sites in AT2 cells with increased age as a disease-relevant epigenetic signature likely controlling basal cell metaplasia.

### *In vitro* IL-1β treatment recapitulates the pro-inflammatory state of AT2 cells from aged lungs and leads to de-repression of AT2 cell metaplasia via a JMJD3-dependent mechanism

We next asked whether age-associated differences in metaplastic potential were reflected in the baseline transcriptional state of freshly isolated human AT2 cells. As previously reported^53–57^, multiple single-cell RNA-sequencing datasets comparing non-diseased human donor AT2 cells from individuals at the extremes of age revealed that the dominant age-associated transcriptional shift is a robust inflammatory transcriptional program, driven largely by NF-κB–dependent signaling. Consistent with these reports, analysis of AT2-cell-specific scRNA-seq datasets generated from 4 non-diseased donors (2 donors <35 years, 2 donors >60 years) revealed a similar age-linked increase in pro-inflammatory gene expression in older AT2 cells (Figure 3A, B, C, D, Supplementary Figure 3). Ingenuity Pathway Analysis (IPA) identified inflammatory signaling as a principal axis of activation with age, with IL-1β–dependent signaling emerging as one of the most prominently activated upstream pathways in older donors. Together, these scRNA-seq data indicate that advanced age is associated with a heightened inflammatory state in AT2 cells, including strong IL-1β pathway activation. To establish a mechanistic link between this inflammatory transcriptional phenotype and the epigenetic changes we observed, we next investigated whether the most important H3K27me3 demethylase in the human genome, JMJD3 (also known as *KDM6B*)^58–61^, was important in de-repression of the basal cell transcriptional program, as several reports have highlighted its role as an important downstream effector of NF-κB signaling in human macrophages^62–64^, as well as a critical regulator of differentiation in embryonic^65^, mesenchymal^66^, and neural stem cells^67^. We functionally tested the roles of IL-1β and JMJD3 during in vitro pre-treatment of AT2 cells from younger donors with IL-1β, with and without a specific JMJD3 antagonist, GSKJ4, prior to trans-differentiation under both mesenchyme co-culture or mesenchyme-free conditions (Figure 3E). For this purpose, we devised a mesenchyme-free trans-differentiation model for AT2 cell metaplasia that relies on roxadustat, a HIF agonist, to drive AT2 cell metaplasia, as others have reported^32,33^. In our preliminary testing, HIF1α inhibition was strongly associated with *KRT5* mRNA reduction in the fibroblast co-culture model (Supplementary Figure 1), and both roxadustat and hypoxia appeared to reliably induce *KRT5* expression in mesenchyme-free conditions in IPF-derived AT2 cells (Supplementary Figure 3). Consistent with our hypothesis, JMJD3 expression levels appeared to be higher in the older donors in our dataset, and using a published IPF Cell Atlas data set as reference in a pseudobulk re-analysis^29^, we found that JMJD3 expression levels were also higher in IPF-derived AT2 cells compared to non-diseased controls (Figure 3F). IL-1β induced expected cytokine and inflammatory markers in AT2 cells from young donors in addition to JMJD3 expression (Figure 3D). Our results revealed that IL-1β pre-treatment led to significant upregulation of both *KRT5* and *KRT17* mRNA in 14-day organoid co-culture (Figure 3E). In the roxadustat model, IL-1β pre-treatment was required for expression of *KRT5* mRNA, while expression of *KRT17* mRNA occurred irrespective of IL-1β pre-treatment, although there was a trend towards significance in the IL-1β pre-treated condition. Remarkably, GSKJ4 administration at the same time as IL-1β pre-treatment abrogated its effects, confirming that JMJD3 is a necessary mediator of this priming, presumably via a NF-κB-dependent pathway, although different or redundant signaling pathways activated by IL-1β may converge on this final epigenetic effector.

**Figure 3.**
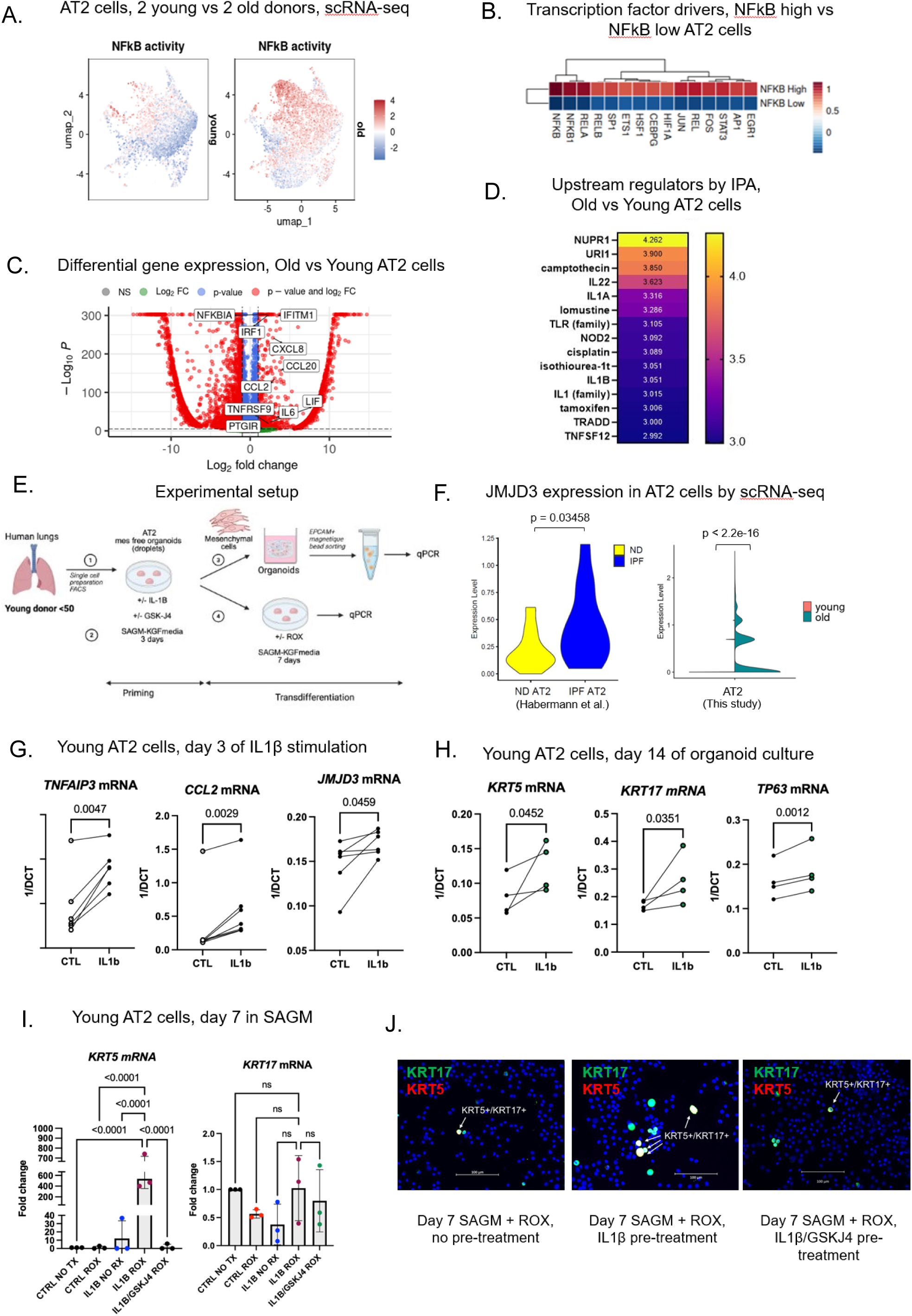
Aged AT2 cells have inflammatory transcriptional changes associated with IL1β and NF-kB activation, and stimulation with IL1β induces priming toward a basal cell fate in young AT2 cells via a JMJD3-dependent mechanism. **(A.)** Single-cell transcriptomic-derived dimension reduction plot (UMAP) of AT2 cells from 2 young (22F, 28M) and 2 old donors (67M, 79F) highlighting higher NF-kB activity scores in old donors, as quantified with decoupleR. **(B.)** Heatmap of transcription factor activity scores between most differentially enriched drivers between NF-kB-high and NF-kB-low AT2 cell clusters across all ages. **(C.)** Volcano plot of differentially expressed genes between old and young AT2 cells across all donors. Highlighted are several inflammatory regulators and cytokines that are more highly expressed in old donors. **(D.)** Heatmap of z-scores from pairwise upstream regulator Ingenuity Pathway Analysis (IPA) of differentially expressed genes between and young AT2 cells. **(E.)** Experimental setup for *in vitro* IL1β stimulation assays. AT2 cells isolated from young donors were treated with IL1β for 3 days and trans-differentiation was then induced either by mesenchymal co-culture or by inducing HIF with roxadustat in mesenchyme-free conditions. **(F.)** Violin plots of scRNA-seq data showing higher expression of JMJD3 in IPF-derived AT2 cells by pseudobulk re-analysis of Habermann et al dataset compared to non-diseased control. Similarly, JMJD3 has higher expression in AT2 cells from older donors in our dataset. Significance testing by two-tailed Wilcoxon rank sum test for single cell data, and one-tailed for pseudobulk data. **(G.)** IL1β stimulation of AT2 cells leads to expression of NF-kB markers such as *TNFAIP3*, chemokines such as *CCL2*, and histone demethylase *JMJD3* by qPCR. **(H.)** IL1β-pre-treated AT2 cells have higher expression of *KRT5*, *KRT17*, and *TP63* in 14-day organoid co-culture with primary fibroblasts by qPCR compared to untreated control. **(I.)** IL1β-pre-treated AT2 cells have higher expression of *KRT5* but not *KRT17* compared to untreated control after 7 days of culture in mesenchyme-free conditions with roxadustat. This effect is abrogated if IL1β pre-treatment is performed with JMJD3 inhibitor GSKJ4. **(J.)** Immunofluorescence staining of cytospins confirming increased KRT5+/KRT17+ basal cells with IL1β pre-treatment of AT2 cells followed by 7 days of mesenchyme-free culture with roxadustat. Both untreated control and IL1β/GSKJ4 pre-treated AT2 cells have <1% KRT5+/KRT17+ basal cells by immunostaining after roxadustat for 7 days, while the number of KRT5+/KRT17+ basal cells is around 5-10% in IL1β pre-treated cells, pointing to a subset of AT2 cells being IL1β-responsive. Statistical significance was determined by unpaired t-test **(F)**, paired t-test **(G, H)**, and ordinary one-way ANOVA **(I)**.

### HIF1α activation redirects KLF5 from AT1-specific to basal-specific binding sites and antagonizes both basaloid-associated SMAD2/3 and AT1-associated YAP signaling

A critical unanswered question from the studies reported thus far is the reconciliation of the seemingly permissive epigenetic state of the *AGER* promoter with the age-dependent decrease in RAGE+ colonies in the organoid co-culture model, as detailed in Figure 1. To better understand the molecular mechanisms leading to increased basal cell trans-differentiation to the detriment of AT1 trans-differentiation in AT2 cells, we focused on KLF5, an essential zinc-finger transcription factor involved in epithelial development, differentiation, and regeneration that was recently implicated in HIF-dependent AT2-to-basal metaplasia^33^. Since KLF5 has been reported to be involved in expression of both KRT5^68,69^ and RAGE^70,71^, we tested whether this transcription factor had a dual role^72^ in AT2 cell fate specification via competitive binding to either basal cell or AT1 genes, in a manner analogous to what has been proposed for NKX2-1^73^. We performed CUT&Tag for KLF5 in AT2 cells isolated from older donors that were treated with LATS inhibitor (LATSI) for three days, which promotes AT2-to-AT1 trans-differentiation and potent RAGE expression via YAP/TAZ signalling^74^, irrespective of donor age. After sampling a portion of the cells for CUT&Tag sequencing, we treated the remaining cells with roxadustat for two additional days, at which point all cells were similarly harvested for sequencing. We also generated KLF5 CUT&Tag sequencing data from freshly isolated IPF-derived basal cells for comparison (see Figure 4A for experimental setup). We confirmed that KLF5 binds to the RAGE promoter in AT1-like cells derived from AT2 cells via LATS inhibition, while it binds to the *KRT5* promoter in IPF basal cells. Importantly, HIF activation via roxadustat treatment leads to decrease in KLF5 binding at the *RAGE* promoter, while its binding was increased at *KRT5*, suggesting re-distribution of its genome occupancy to favor a basal cell fate (Figure 4B). Consistent with this, there is a decrease in KLF5 enrichment around the transcription start sites of AT1-associated genes with roxadustat, and even more so in IPF-derived basal cells (Figure 4C), while the proportion of genes bound by KLF5 that are shared with IPF basal cells increases slightly (77% to 81%) after roxadustat treatment in AT1-like cells (Figure 4D). This implicates the epigenetic state of basal cell-associated KLF5 binding sites as a key factor controlling whether the AT2-to-AT1 transcriptional program can be activated in the presence of hypoxia or HIF stabilization. To further elucidate regulation of KRT17 and explain discrepancies with KRT5 expression, despite having similar age-dependent de-repression, we performed mesenchyme-free induction of KRT5-/KRT17+ basaloid fate in AT2 cells from older donors with TGFβ1 treatment. We then tested the role of JMJD3 in this pathway given its higher expression in basaloid cells by again using concomitant JMJD3 inhibition with GSKJ4 (Figure 4E and F, Supplementary Figure 4). JMJD3 activity was necessary for *KRT17* mRNA expression, suggesting direct recruitment of this effector to downstream genes by the TGFβ1-SMAD signaling axis. Consistent with this finding, *KRT17* mRNA expression in response to TGFβ1 was not age-dependent, suggesting a reduced role for priming and differential epigenetic regulation compared to HIF-dependent activation of *KRT5* (Figure 4G). While HIF also induces *KRT17*, as demonstrated by KLF5 binding to both promoters, our experimental data suggests that the effects of TGFβ1 activity on *KRT17* expression may be stronger.

**Figure 4.**
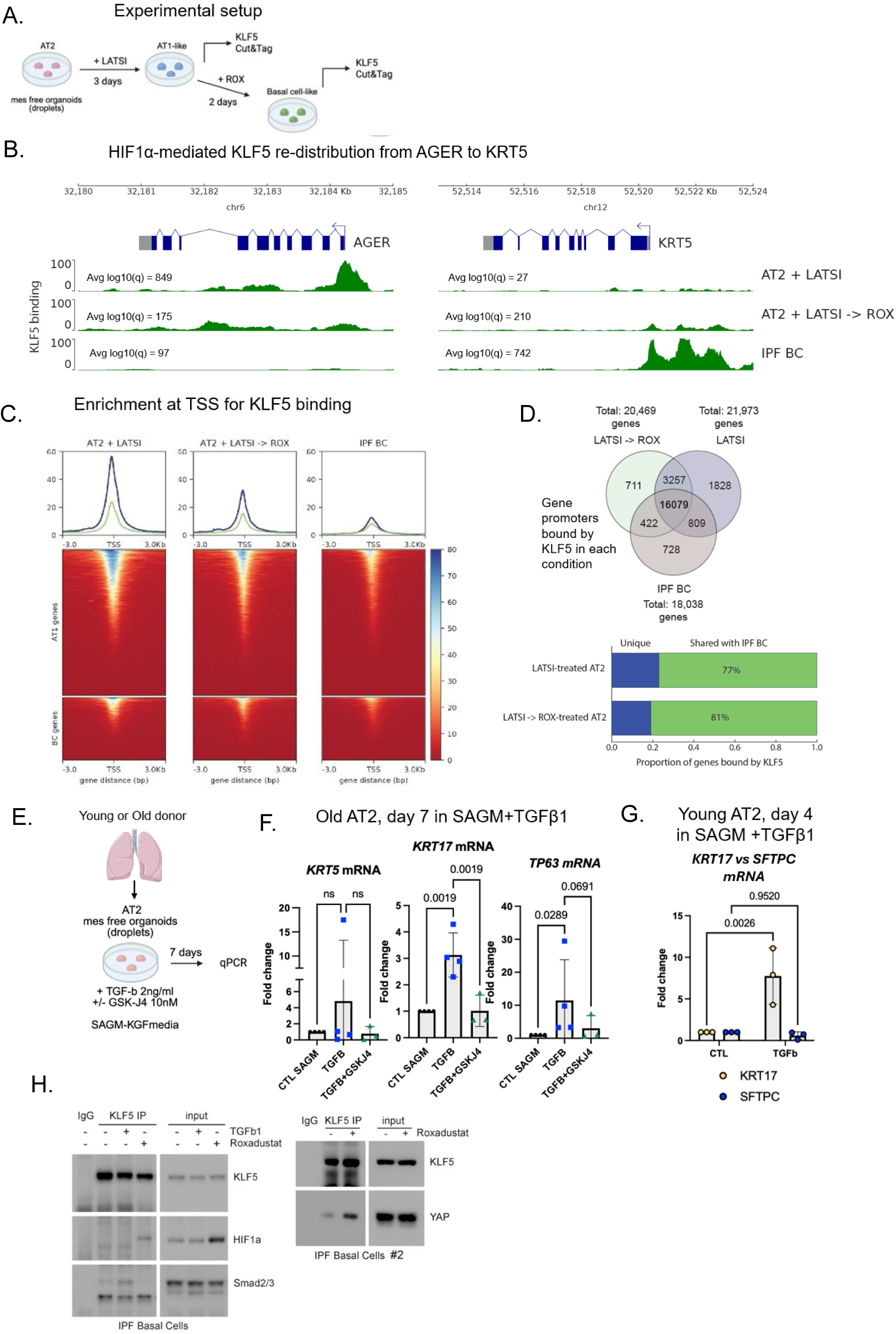
HIF activation leads to KLF5 re-distribution from AT1-associated loci to basal cell-associated loci in old AT2 cells and antagonizes KRT17 activation by the TGFβ-SMAD2/3 signaling axis, which is specific to a basaloid cell fate. **(A)** Schematic of experimental setup for CUT&Tag profiling of KLF5 binding sites. AT2 cells from old donors were treated with LATS inhibitor to induce AT1 marker expression via YAP/TAZ activation for 3 days, a portion of cells harvested for CUT&Tag, and the rest cultured in the presence of HIF agonist roxadustat for 2 days to drive acquisition of basal cell markers, prior to being collected for repeat CUT&Tag. Results were compared with CUT&Tag for KLF5 in NGFR+ basal cells from an IPF donor. **(B)** Representative genomic tracks of normalized read depth from CUT&Tag sequencing for KLF5 in AT2 cells treated with LATS inhibitor for 3 days to drive an AT1-like cell state (track 1, “AT2 + LATSI”), treated with LATS inhibitor for 3 days and then roxadustat for 2 days (track 2, “AT2 + LATSI -> ROX”), and finally IPF-derived basal cells (track 3, “IPF BC”). Select marker loci (*AGER* as AT1 cell marker, *KRT5* as a basal cell marker) are shown. Peak height represents stronger KLF5 binding, while −log10(q) values shown represent the average statistical significance of relevant peak calls in the promoter region. There is strong binding to *AGER* promoter with LATS inhibition that decreases with HIF agonism, while *KRT5* promoter binding increases. Conversely, there is strong binding to *KRT5* promoter in IPF-derived basal cells and no significant binding to *AGER* promoter. Of note, KLF5 also binds the *KRT17* promoter (not shown) in both roxadustat treated and IPF basal cells. **(C)** Enrichment around transcription start site (TSS) for KLF5 in differentially bound genes between LATS-inhibitor treated cells and IPF-derived basal cells. Genes with higher promoter binding in each condition were labeled “AT1 genes” and “BC genes”, respectively. **(D)** Identification of shared genes with any KLF5 binding (peak q-value cutoff < 0.01) within their promoter region (defined as TSS +/− 3kb) across conditions, irrespective of peak height (i.e. binding intensity). After treatment with roxadustat, a greater proportion of KLF5-bound genes are shared with IPF basal cells in LATS-treated AT1-like cells (increase from 77% to 81%). **(E)** Schematic of experimental setup for basaloid cell fate induction in mesenchyme-free conditions in old AT2 cells. **(F)** Activation of *KRT17* by TGFβ1 is JMJD3-dependent and is specific for the basaloid cell fate (KRT5- /KRT17+). **(G)** Expression of *KRT17* does not depend on AT2 cell priming or age, as shown by robust mRNA expression in young AT2 cells by day 4 of TGFβ1 treatment. **(H)** Immunoprecipitation of KLF5 and Western Blot for SMAD2/3, HIF1α, and YAP from IPF-derived basal cells treated with either roxadustat or TGFβ1. HIF1α stabilization with roxadustat drives its association with KLF5 while also diminishing KLF5-SMAD2/3 association, which instead is strongest in the presence of TGFβ1. Similarly, HIF1α stabilization also drives association of KLF5 with YAP, a key driver of AT2-to-AT1 transdifferentiation. Statistical significance was determined by Poisson local test with BH correction as in Figure 2 **(A)**, by ordinary one-way ANOVA **(F)** and two-way ANOVA **(G)**.

Given that KLF5 is reported to also physically interact with SMAD2/3^75,76^, a well characterized target of TGFβ1 signaling, we sought to dissect the role this key transcription factor plays in basal, basaloid, and AT1 cell trans-differentiation with biochemical methods. We performed immunoprecipitation of KLF5 in IPF-derived basal cells in the presence of HIF1α stabilization by roxadustat treatment, or SMAD2/3 activation by TGFβ1 treatment, after in vitro culture for 12 hours, followed by Western blotting for HIF1α, YAP, and SMAD2/3 (Figure 4H). Compellingly, we found that in the presence of HIF1α stabilization by roxadustat, KLF5 complexes with both HIF1α and YAP, suggesting that suppression of AT1 cell fate may be due to direct recruitment of KLF5 and YAP by HIF1α to de-repressed p63 target sites in AT2 cells from older donors. We were unable to detect p63 by Western blot in IPF basal cells, presumably due to low expression levels, but we separately confirmed that p63 also co-precipitates with KLF5 in a squamous cancer cell line, CAL27 (Supplementary Figure 4). Consistent with this hypothesis, treatment with TGFβ1 leads to increased association between KLF5 and SMAD2/3 compared to untreated control, while HIF1α activation antagonizes this association to favor HIF1α-KLF5 complexing instead. Taken together, these findings suggest that there are opposing molecular mechanisms governing AT2 cell fate choice, and that KLF5 is a central regulatory transcription factor that limits the differentiation trajectories based on its active interacting protein partners and the epigenetic state of the genome.

## Discussion

Our study highlights important phenotypic and epigenetic differences in primary human AT2 cells across the human lifespan that provide a mechanistic basis for the increased risk with age for IPF. Collectively, these findings outline a model where chronic inflammation, one of the hallmark features of tissue and organismal aging, may directly lead to epigenetic changes in AT2 cells, and possibly other stem cell niches. These epigenetic changes in turn affect AT2 differentiation potential, and result not only in stem cell exhaustion, but also acquisition of metaplastic lineage potential, with important implications for tissue regeneration. Importantly, aging-associated epigenetic de-repression hinges on partial removal of a repressive histone mark, H3K27me3, to allow for deposition of H3K4me3 and establishment of bivalency, as the two marks cannot co-exist on the same histone tail^77,78^. This does not result in transcriptional activation, but rather in a re-programming of the “priming” status of key genes that results in increased AT2 susceptibility to chronic stress and injury, which is ultimately the trigger for the alveolar epithelial remodeling seen in IPF. We further identify important distinctions in the regulation of KRT5-/KRT17+ basaloid and KRT5+/KRT17+ basal cell fate by TGFβ1 signaling or HIF1α activation, respectively, which may provide valuable clues as to their evolutionary function in lung injury and fibrosis.

Trans-differentiation into aberrant lineages requires specific signals, such as hypoxia, a powerful stressor that involves HIF stabilization and activation. Our data suggests that HIF1α activity is both sufficient and necessary to induce expression of basal markers in AT2 cells. We define a clear molecular mechanism that hinges on KLF5 re-purposing from a driver of AT1 cell differentiation to a driver of metaplasia when HIF1α is active. This re-purposing is driven by formation of a transcription factor complex when stable HIF1α is present that includes YAP, a key master regulator of AT1 cell fate. Shared transcription factors such as KLF5 and YAP, needed for activation of the AT1 transcriptional program, are thus shunted to alternate genomic binding sites by participation in this complex. Although p63 protein could not be detected due to low amounts in either alveolar epithelial cells or even basal cells, using a squamous cell carcinoma line we found that p63 is recruited as part of this complex and likely acts as a crucial DNA binding component that directs the HIF1α-KLF5-YAP complex to poised p63 binding sites in aged AT2 cells, such as the KRT5 promoter. While we cannot exclude the simultaneous participation of KLF5 in separate complexes, such as KLF5-YAP and KLF-HIF1α, the shifts in genomic occupancy observed under HIF agonism by direct CUT&Tag profiling of KLF5 DNA binding activity are compelling. The lack of significant regulation at the level of histone marks or chromatin accessibility at AT1 marker genes such as AGER suggests that this re-distribution of transcriptional machinery is mediated by epigenetic de-repression of basal cell genes in primed AT2 cells and recruitment to these sites by active HIF1α signaling, rather than epigenetic repression of the AT1 transcriptional program.

The exact triggers and chronicity of stimuli that account for the establishment of a de-repressed epigenetic state in human AT2 cells at p63 binding sites over the lifetime of an individual remain to be defined. Our data supports the presence of local tissue inflammation in aged lungs, possibly via secretion of IL-1β by alveolar macrophage^79,80^ or tertiary lymphoid structures, such as bronchus associated lymphoid tissue^81,82^ (BALT), which may develop over many years of repeated mild to moderate injury, such as aspiration or viral infection. These paracrine signals are the likely cause of increased pro-inflammatory markers, including NF-KB activation, in a subpopulation of AT2 cells that appears to increase in abundance during normal aging. Inflammatory signals, such as IL-1β, induce epigenetic de-repression of AT2 cells from younger donors in a manner reminiscent of what has been described in murine models as IL-1β-mediated priming toward formation of KRT8+ damage-associated transient progenitors (DATPs) in response to hypoxia^83^. While there is no exact human equivalent for this cell population, KRT8+ transitional cells may represent a non-specific activated AT2 cell state, that in the presence of appropriate p63-related priming, could precede trans-differentiation into a basal cell fate.

We posit that IL-1β activates synergistic pathways in AT2 cells, including NF-KB, that converge on JMJD3-mediated histone demethylation and downstream activation of bivalent gene promoters. JMJD3 and the less abundant UTX/UTY act as the main H3K27me3 demethylases in the human genome^58–61^. Given that UTX/UTY are mostly active during embryonic development^84–86^, JMJD3 likely plays a key role in the establishment of promoter bivalency during aging as a specific manifestation of what has been termed inflammatory epigenetic memory^87^. It is likely that additional factors induced by inflammatory signals, such as the AP1 family of transcription factors, interact with these epigenetic modifiers to remove H3K27me3 and allow H3K4me3 deposition by methyltransferases such as MLL2^88^ at a subset of target sites in a coordinated fashion. Additional research is needed to understand how inflammatory stimuli induce specific epigenetic changes in AT2 cells, whether these epigenetic signatures are heritable over multiple mitotic cycles even in the absence of ongoing inflammatory stimulation, or whether maintenance of bivalency is predicated on the presence of ongoing paracrine inflammatory mediators, with important implications for whole tissue aging. Our data suggests that these cells retain their primed epigenetic state during short term *in vitro* culture.

Lastly, while de-repression of the basal cell transcriptional program seems to require prior IL1β and JMJD3 induction in a “two-hit” model (Figure 5), expression of KRT17 by the well-described TGFβ1-SMAD2/3 signaling axis is not influenced by age or epigenetic status, suggesting that TGFβ1 may directly recruit epigenetic remodelers such as JMJD3 to ensure transcription of its downstream targets^89,90^. Indeed, HIF activation seems to antagonize SMAD2/3-KLF5 complex formation, and favor HIF1α-KLF5 formation, pointing to a role for KLF5 in regulating alternate transcriptional programs. While single cell transcriptomic maps place basaloid cells as likely intermediates between AT2 and basal cells, these approaches have important limitations^91^. The lack of age-dependence may hint at an adaptive role for the KRT5-/KRT17+ basaloid cell population in lung injury. Additional functional characterization of these aberrant cell populations and their lineage trajectories is needed to fully understand AT2-specific contributions to epithelial remodeling in lung fibrosis. It is important to note that mesenchymal cells, while not contributing to the aging-associated phenotype we observed in our *in vitro* model, play an important role in controlling basal and basaloid cell formation in IPF, including by modulating Wnt signaling in AT2 cells^92,93^. Given the high degree of tissue complexity in fibrotic lungs, and the presence of multiple cell type compartments (epithelial, immune, mesenchymal, and endothelial), additional signals likely shape AT2 cell trans-differentiation capacity other than what we describe in our study. A better understanding of how integration of these differentiation signals occurs in AT2 cells and their interaction with the epigenetic status of the cell, especially in terms of how chromatin remodelers such as JMJD3 are affected, is going to be critical to manipulate this system to favor reparative pathways promoting AT1 and AT2 regeneration.

**Figure 5.**
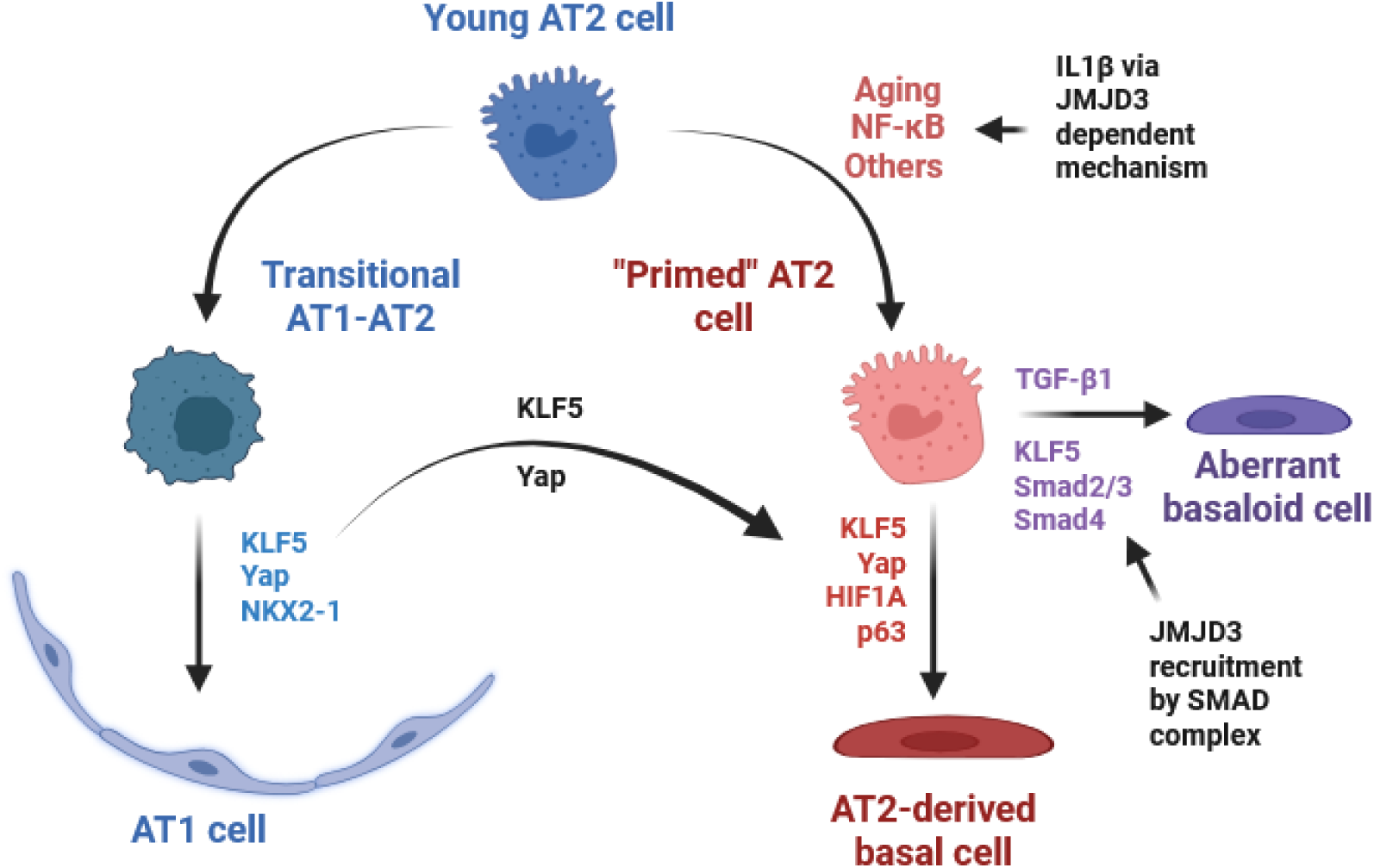
Graphical summary.

## Methods

### Sex as a biological variable

Sex was not considered as a biological variable. Lung donors of both sexes were used in this study.

### Human lung tissue processing and fluorescence activated cell sorting (FACS) of epithelial cell types of interest

Lungs from non-diseased donors declined for clinical use in transplantation, excess non-diseased tissue from surgical biopsies, and explanted lungs from IPF patients undergoing transplantation were de-identified and donated to research through institutional protocols approved by the UCSF Institutional Review Board (Supplementary Table 1). A single cell preparation was prepared from these tissues by mechanical disruption and enzymatic digestion (dispase,15 I.U./mL and collagenase, 225µg/mL), as previously described (Ref). FACS was performed on digested donor lung tissue with to obtain AT2 cells (EPCAM^pos^/CD11b^neg^/CD31^neg^/ CD45^neg^/ HTII-280^pos)^, mesenchymal cells (EPCAM ^neg^/CD11b^neg^/CD31^neg^/CD45 ^neg^), and basal cells (EPCAM^pos^/CD11b^neg^/CD31^neg^/ CD45^neg^/ HTII-280^neg^/NGFR^pos^). For IPF tissues, HTII-280 positivity in the AT2 cell population appeared to be spread out over a larger range, so our gating strategy focused on sorting only HTII-280^high^ AT2 cells. Comparison with a EPCAM^pos^/CD11b^neg^/CD31^neg^/ CD45^neg^/ HTII-280^pos^/NGFR^neg^ gating strategy yielded the same population with >99% overlap, and immunofluorescence staining of cytospins (see relevant Methods section below) confirmed selectivity for AT2 cells with either sorting strategy in IPF donors. The list of antibodies used is provided in Supplementary Table 2.

### In vitro and ex vitro models

1) *Cell culture*. Adult human mesenchymal cells were cultured in DMEM (Cat#11965092, Thermo Fisher) with 10% fetal bovine serum (Cat#SH883IH2540, Fisher Scientific), 1% GlutaMAX (Cat#35050-61, Gibco), 1% HEPES (Cat#5630-080, Gibco), and 1% Pen/Strep (Cat#10378016, Gibco). Cells were used within the first five passages of being isolated from donor lungs for mesenchymal cells. Sorted AT2 were cultured in 3D conditions embedded in Matrigel. For each droplet, 5,000 to 10,000 AT2 cells in 20 µl of SFFF media supplemented with 2% human serum diluted 1:1 in growth factor-reduced Matrigel (Cat# CB-40230A, Thermo Fisher) were deposited onto the bottom of a 12-well culture plate (3 droplets/well) as previously described^94^. Media was changed every 2-3 days. When applicable, cells were treated with IL-1β (25ng/ml, HY-P7028, Med Chem Express), GSKJ4, a JMJD3 inhibitor, (5-10μM, HY-15648B, Med ChemExpress), BAY 11-7082, an NF-KB inhibitor, (10 –25 µM, Cat# 196870, Sigma), roxadustat, a prolyl hydroxylase inhibitor and HIF agonist (15 μM, FG-4592, MedChemExpress), TGFβ1 (2ng/mL, Cat#100B, R&D Systems), or LATS inhibitor (LATSI or Lats-IN-1, 10uM, HY-138489, MedChemExpress). Prior to treatment, media was changed for SAGM-KGF supplemented with 5% charcoal treated FBS and 1% Pen/Strep. Where applicable, the effects of hypoxia were tested in AT2 medium, and by placing cells in an airtight chamber in 2% oxygen for 3 days without any media changes. 2) *Organoid assays*. AT2 cells and mesenchymal cells were co-cultured (5,000 AT2:30,000 fibroblasts/mesenchymal cells per well) in modified MTEC medium diluted 1:1 in growth factor-reduced Matrigel (Cat# CB-40230A, Thermo Fisher). The cell suspension–Matrigel mixture was placed in a transwell and incubated with 10 μM Y-27632 ROCK-inhibitor compound (Cat# 1254, Tocris) for the first 24 hours. Isolation of the cells from organoids was performed as described below. Each experimental condition was performed at least in triplicates or as indicated in figure legends. Where applicable, HIF inhibitors, PX-478 (HIF1α-specific inhibitor, HY-10231, MedChemExpress) and PT-2385 (HIF2α-specific inhibitor, HY-12867, MedChemExpress) were added to the media at a final concentration of 20 μM and replenished in every medium change for days 7-21. Organoids were processed for OCT-embedding for immunostaining or made into a single cell preparation and positively selected via magnetic beads for EPCAM+ prior to RNA extraction. The composition of all media (SFFF, MTEC, SAGM-KGF) is reported in Supplementary Table 3.

### Single cell preparation from 3D organoids and droplets

1) *Organoids*. The cell–Matrigel mixture in the transwell was washed with PBS and incubated in the 15 µg/mL dispase for 30 minutes at 37°C with intermittent resuspension. The mixture was removed from the transwell and resuspended in TrypLE (Cat# 12563011, ThermoFisher). Cells were shaken at 37 °C for up to 20 minutes, pipetting up and down 10 times every 5 minutes and checking for single cells.

For organoid isolation only, cells were then incubated with biotin anti-CD326 (Cat# 324216, BioLegend) for 30 minutes at 4°C. Streptavidin beads (Cat# 17663, STEMCELL) were added to isolate the epithelial cells, while the rest of the unbound cells consisted of mesenchymal cells. 2) *Droplets*. AT2 cells cultured in mesenchyme-free 3D Matrigel droplets were similarly washed twice with PBS, dispase (5U/mL) was added, and plates were incubated for 35 minutes shaking at 37°C. For single cell suspension, the dissolved cell-Matrigel mixture was incubated with TrypLE as above. Wells were washed twice with PBS to ensure recovery of all cells.

### Immunofluorescence and quantification

1) *Optimal Cutting Temperature (OCT) embedding.* Organoids were fixed with 4% PFA (Cat# 15714, Electron Microscopy Sciences), and embedded in OCT after undergoing a sucrose gradient, as previously described^25^. Sections (7μm in thickness) were cut on a cryostat. 2) *Cytospins of sorted cells*. Cell pellets were fixed in 4% PFA then loaded into chambers and spun onto Superfrost Plus microscope slides (Cat# 12-550-15, ThermoFisher). Slides were subsequently immunostained. 3) *Immunofuorescence staining.* Prior to antigen retrieval, OCT-embedded slides were fixed in 4% PFA. Antigen retrieval (Cat# DV2004MX, Biocare) was performed. Slides were washed with PBS, blocked/permeabilized (5% horse serum 0.5% BSA 0.1% Triton X) for 1 hour, and then incubated with primary antibodies overnight at 4°C and with Alexa Fluor secondary antibodies for 1 hour (see Supplementary Table 2 for list of antibodies). Images were captured using ZEN v3.1 software (Zeiss); 3) *Image quantification.* Slides were imaged for quantification on a Zeiss AxioImager.M1 microscope. Cell counts for stained organoids or cytospins were performed manually. Approximatively 50 organoids or 1,000 cells (for cytospins) per condition were counted blindly by members of the laboratory. The results were averaged between each specimen and S.D. values were calculated for each condition. The organoids were accrued over a period of 18 months and stored uncut in OCT, at which point they underwent immunofluorescence staining for AT2, AT1, and basal cell markers as described in the main text and evaluated as a batch by two observers blinded to donor identity and age. A colony was determined to be positive for KRT5 or RAGE if at least 25% of the cells were positive.

### RNA extraction and Quantitative RT-PCR

RNA was extracted using the ReliaPrep RNA Cell Miniprep System (Cat# Z6011, Promega) as per manufacturer’s instructions. Reverse transcription was performed with iScript RT Supermix (Cat# 1708841, Bio-Rad) and quantitative real-time PCR (qPCR) was performed using Sso Advanced Univ SYBR Green Supermix (Cat# 1725271, Biorad). Relative mRNA expression levels were calculated with the delta-delta method. The list of primers is provided in Supplementary Table 4.

### Immunoprecipitation and Western blotting

1) *Immunoprecipitation*. Basal cells isolated from IPF donor lungs via FACS were grown on collagen coated plates in Pneumocult medium (Supplementary Table 3). Culture medium was supplemented with 10 μM Y-27632 ROCK-inhibitor for 24 hours after each passage (Cat# 1254, Tocris). The CAL-27 cell line was cultured on plastic in DMEM, supplemented as for fibroblast culture. Cells were lysed in NP40 lysis lysis buffer (50 mm Tris-HCl, pH 7.5, 150 mm NaCl, and 1% NP40) supplemented with protease inhibitors and 1 mM phenylmethylsulfonyl fluoride. Clarified lysates were immuno-precipitated with antibody to KLF5. The precipitates were blotted for KLF5, HIF1α, SMAD2/3, p63, or YAP. 2) *Western blot.* Cells were lysed in RIPA buffer (150 mM NaCl, 50 mM Tris, pH 8.0, 1% Triton X-100, 0.5% sodium deoxycholate, 0.1% SDS supplemented with protease inhibitor, and 1 mM phenylmethylsulfonyl fluoride). Equal amounts of protein were loaded per lane and separated by SDS-PAGE. The protein was transferred to nitrocellulose membrane and blotted for different proteins. The same membrane was blotted for β-actin as loading control.

### Bulk ATAC-seq analysis

For bulk ATAC-seq, approximately 100k-200k freshly isolated AT2 cells from IPF donors (n=3) and age-matched controls (n=3) (see Supplementary Table 1 for donor information) were processed using standard Active Motif manufacturer’s protocols. The resulting FASTQ files were inspected for quality control with FastQC and mapped against the GRCh38 reference genome using bwa-mem. Duplicate reads were removed with Picard. Peaks were called and quantified with MACS3 callpeak function. Differential peak analysis was performed using DiffBind (v3.23) in R (v4.3.1). Gene set enrichment analysis was performed using the manually curated ChEA3 database (Cit), which integrates transcription factor vs target gene association libraries from ChIP-seq experiments from ENCODE, ReMap, and additional individual publication; co-expression data from GTEx and ARCHS4; co-occurrence data from Enrichr; and transcriptional gene signatures from single transcription factor perturbation experiments from additional individual publications.

### Histone and KLF5 CUT&Tag analysis

For histone CUT&Tag, approximately 200k freshly isolated AT2 cells from donors of different ages (Supplementary Table 1) were processed with CUT&Tag kit following standard manufacturer’s recommendations. For KLF5 CUT&Tag, AT2 cells were cultured in the presence of LATS inhibitor (LATSI) for 3 days or LATS inhibitor for 3 days followed by roxadustat for 2 days as described in the main text. Cells were then harvested and made into a single cell suspension before being processed with Active Motif CUT&Tag kit using manufacturer’s protocol, using approximately 200k cells as input. Given the fragility of LATSI or roxadustat-treated primary AT2 cells, we opted for a lower digitonin concentration (2.5%) for overnight permeabilization and used 2uL of primary KLF5 antibody. Separately, we isolated basal cells from IPF tissues via FACS and processed the resulting single cell suspension following standard Active Motif CUT&Tag protocols. See separate antibody table for information on KLF5, H3K4me3, and H3K27me3 primary antibodies (Supplementary Table 2). A negative IgG control, which did not yield a cDNA library, and a positive control using H3K4me3 were included, but not sequenced. Sequencing was performed on the Illumina Novaseq platform. Resulting FASTQ files were mapped against GRCh38 human reference genome after quality control with FastQC. Mapping was done using bowtie2 with end-to-end, very-sensitive, and no-discordant flags, retaining only properly paired mapped reads. Duplicate reads were marked with Picard but not filtered out, given that reads with exact start and end coordinates are common at modified Tn5 insertion sites in CUT&Tag data. Peaks were called with MACS3 callpeak function, using a q score cutoff of 0.01 for KLF5, and a cutoff of 0.1 for histone marks. For H3K27me3, the “broadpeak” option was used instead of “narrowpeak”, as these histone marks are known to cover longer regions of chromatin compared to most transcription factor binding sites and H3K4me3 histone marks. Consensus peak calls in triplicates was performed as detailed in Supplementary Figure 2 for histone marks. The resulting bedgraph/bigwig files were processed with bedtools, deeptools, and pygenometracks for plotting and interpretation.

### Single-cell RNA analysis

FASTQ files generated through 10x Genomics standard 3’ gene expression scRNA-seq platform followed by Illumina Novaseq sequencing in freshly sorted EPCAM+ epithelial cells from two donors (79F, 28M) were run through CellRanger (v8.0.0) software with default settings for de-multiplexing, aligning reads with built-in STAR algorithm to hg19 or GRCh38, and counting unique molecular identifiers (UMIs). Seurat package (v5.3.0) in R (v4.3.1) was used for downstream analysis. Single-cell transcriptomes were also obtained from GEO (GSM3489193)^95^ and processed using Seurat. Low-quality cells were filtered (>15% mitochondrial reads). Principal component analysis was performed using SCT-transformed normalized and integrated data. The top 30 principal component analyses were inputted in the UMAP algorithm from the Seurat package for visualization and clustering. Annotation was done using the Azimuth package (v0.5.0) AT2 cells were extracted, re-clustered, and analyzed. The lists of DEGs were identified with a non-parametric Wilcoxon rank sum test and filtered (adjusted p-value<0.05 and percent of cells expressing gene>10%). Transcription factor activity analysis was done using the decoupleR package (v2.9.7). Qiagen Ingenuity Pathway Analysis (IPA) was performed on significant differentially expressed genes across conditions to identify upstream regulators. Where applicable, data from IPF Cell Atlas (GEO Accession Number: GSE135893)^29^ was sub-setted for AT2 cells, basal cells, and basaloid cells from IPF and healthy donors, and pseudo-bulked by donor using Seurat prior to comparison of expression levels of candidate genes by Wilcoxon rank sum test.

### Statistics

Statistical analyses for cell count and gene expression were performed in GraphPad Prism. The two-tailed Mann-Whitney test was used for statistical analysis of scRNA-seq gene signature expression. One-Way ANOVA, unpaired and paired two-tailed t-tests were used to determine the P values, and the data in the graphs are presented as mean ± S.D. Unpaired t-test was used to compare two treatment groups. The Kruskal–Wallis test or Dunnett’s multiple comparisons test were used for multiple comparisons. For normally distributed data, ordinary one-way ANOVA followed by Tukey’s multiple comparisons test was performed.

## Data availability

The bulk ATAC-seq and CUT&Tag data that support the findings in this study have been deposited in the Gene Expression Omnibus (GEO) under the following accession codes (pending upload). Previously published scRNA-seq data that are re-analyzed here are available in GEO (GSM3489193, GSE135893). Additional scRNA-seq data generated for this study is similarly available in GEO (pending upload). All other data supporting the findings of this study are available from the corresponding author on reasonable request. Source data are provided with this paper.

## Code availability

No custom bioinformatic tools were developed or used in this manuscript. Pipelines were manually curated from publicly available Galaxy tutorials and optimized to run on the UCSF Wynton high-performance computing cluster (HPC). All scripts and codes are available by request to the corresponding author.

