## Supplementary Figures 1-4 for "Epigenetic de-repression of basal cell metaplasia in aging AT2 cells is a risk factor for idiopathic pulmonary fibrosis (IPF)"

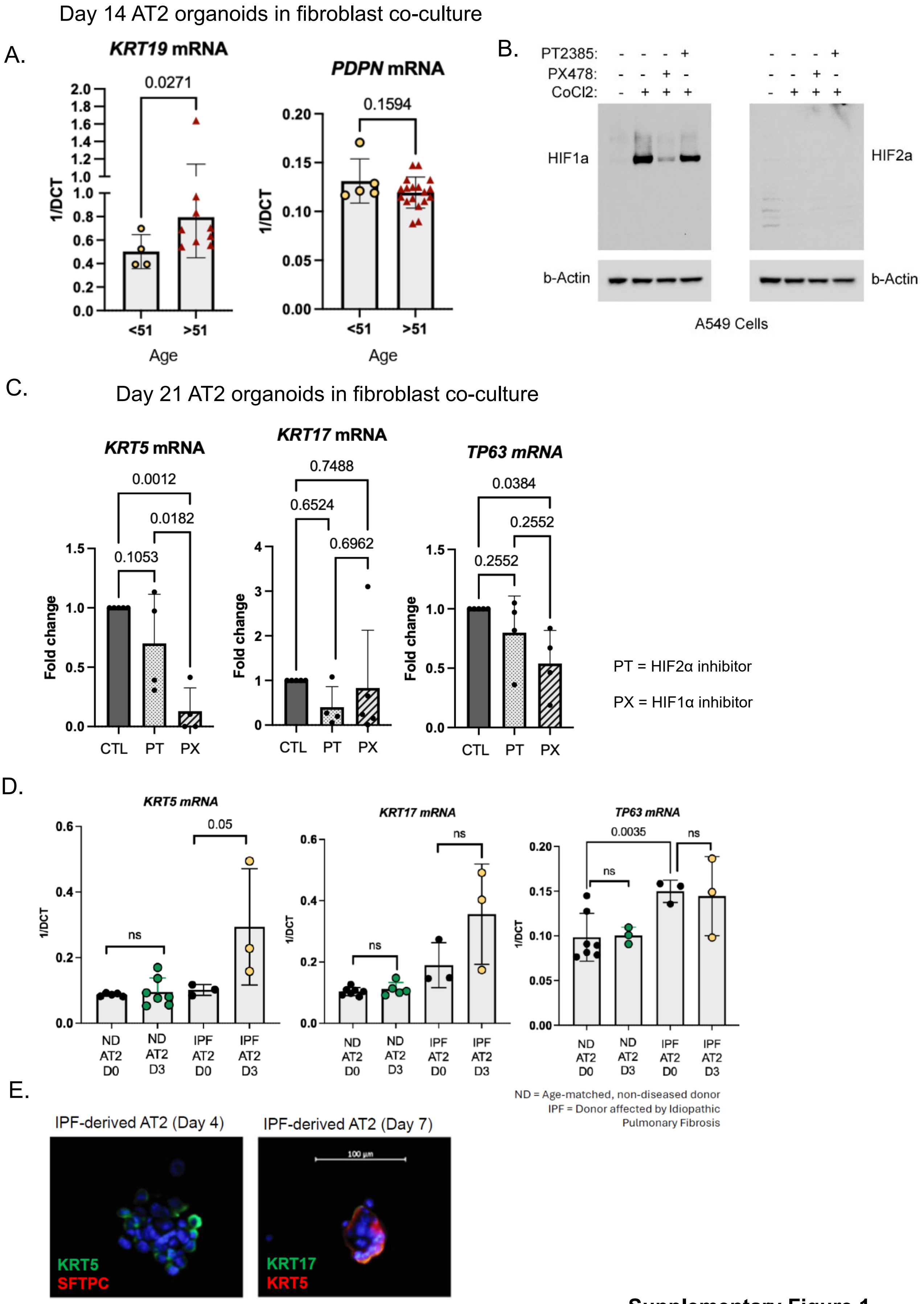


**Supplementary Figure 1**

**Supplementary Figure 1. Additional gene markers in 14-day AT2 organoid co-culture with primary fibroblasts by donor age. (A)** Expression data by qPCR for *KRT19* mRNA, a cytokeratin marker of basal cells, was significantly more increased in organoids derived from donor >51 in age. Instead, *PDPN* mRNA, a marker of AT1 cells, was not significantly different between age groups. **(B)** Validation of HIF1α-specific inhibitor PX478 and HIF2α-specific inhibitor PT2385 in A549 cells treated with cobalt chloride to stabilize HIF proteins. **(C)** Treatment with PX478 (PX) leads to greater reduction in *KRT5* mRNA than treatment with PT2385 (PT), suggesting a greater role for HIF1α in activation of basal cell transcriptional program in AT2 cells trans-differentiating in the fibroblast co-culture organoid system for 21 days. Treatment occurred for 14 days, with 1 initial week allowed for cultures to take hold. **(D)** IPF-derived AT2 cells, isolated by IPF donor lungs using flow cytometry using a HTII-280-high sorting strategy and grown in SAGM-KGF media for 3 days, acquire higher *KRT5* mRNA expression levels by qPCR compared to age-matched non-diseased donors, suggesting a primed state. **(E)** Immunofluorescence staining confirmed presence of KRT5+ and KRT17+ cells at day 4 and day 7, but the fraction was less than 5% of total AT2 cells cultured. As expected SFTPC was mostly lost in AT2 cell cultures by 4 days. Statistical significance was determined by unpaired t-test **(A, D)** and one-way ANOVA **(C)**.

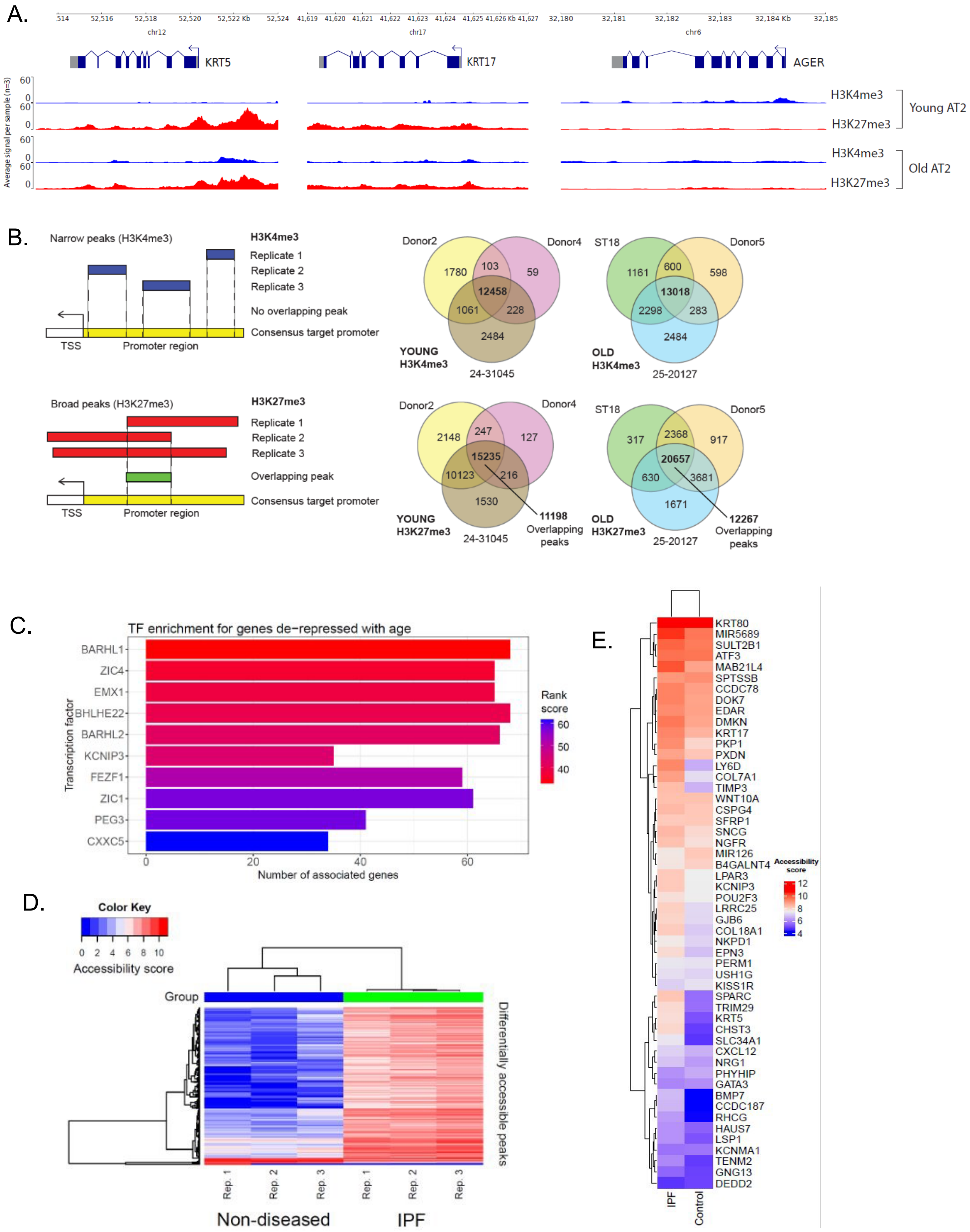


**Supplementary Figure 2**

**Supplementary Figure 2. Histone CUT&Tag for "Young" and "Old" samples and bulk ATAC-seq for IPF and age-matched non-diseased samples. (A)** Representative plots showing average normalized signal across 3 replicates for each condition at *KRT5*, *KRT17*, and *AGER* loci. **(B)** Workflow to identify consensus peaks for H3K4me3, which are typically "narrow" peaks, and H3K27me3, which are "broad" peaks. Peak size affects overlap between replicates. For each condition, we identified promoter regions (TSS +/- 3kb) that were marked by the same histone modification in all 3 replicates. For H3K4me3, we did not perform additional filtering. For H3K27me3, since the peak coordinates tend to overlap with each other across replicates due to their larger size, we further identified overlapping regions in peak calls across replicates, thus reducing the number of gene promoters marked by these "overlapping peaks". **(C)** ChEA3 database gene set enrichment analysis for all de-repressed genes does not yield known epithelial regulators as significantly enriched in top 10 ranked transcription factors. **(D)** Genome-wide accessibility score for significant differentially accessible peaks across 3 replicates for IPF vs age-matched non-diseased comparison, showing that the majority of significant peaks are more "open" in IPF. **(E)** List of p63 target genes that are both de-repressed with age and more accessible in IPF, showing average accessibility per gene in IPF vs age-matched non-diseased.

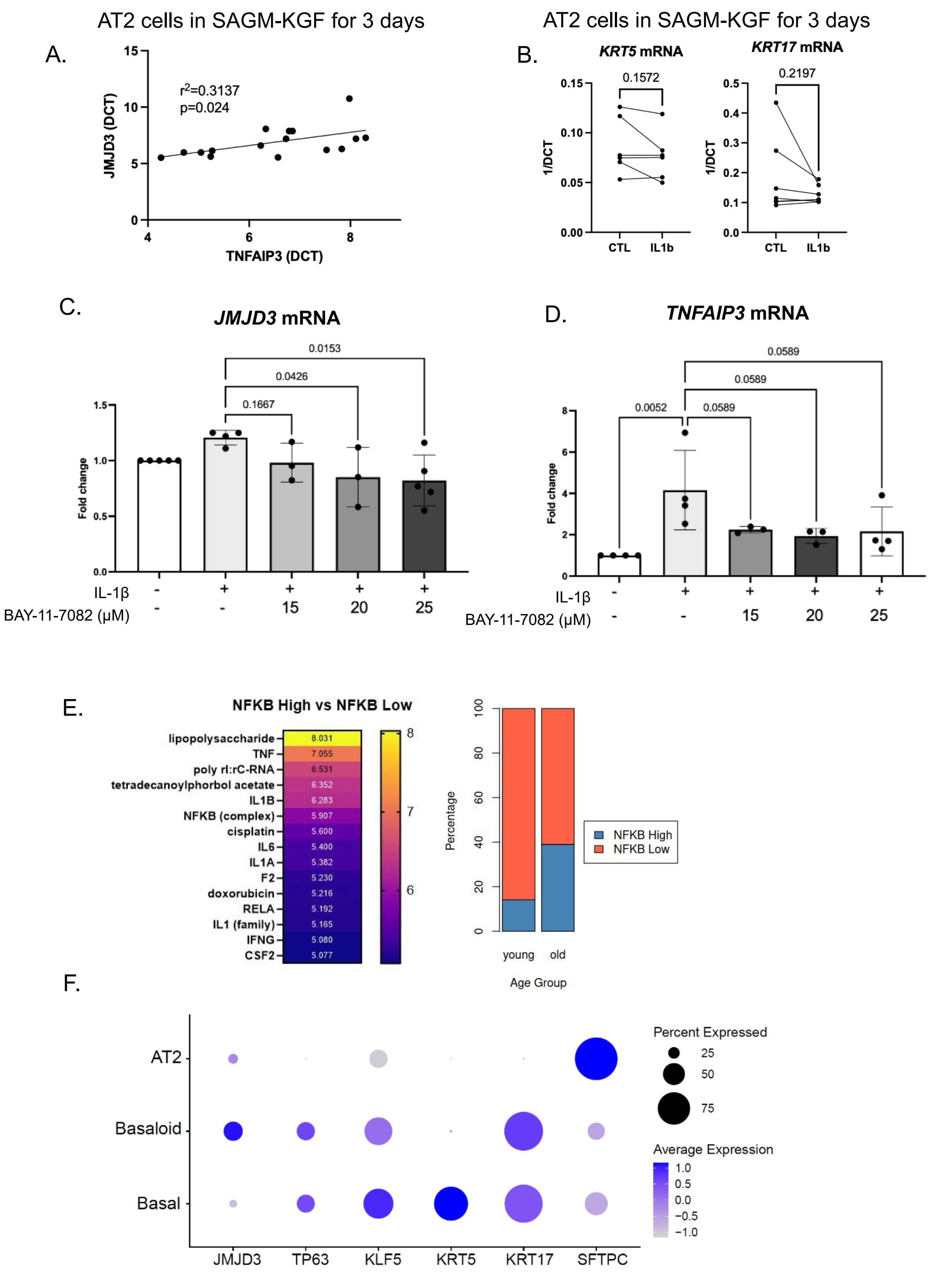


**Supplementary Figure 3**

**Supplementary Figure 3. IL1β stimulation of *in vitro* cultured AT2 cells from "young" donors and scRNA-seq analysis of AT2 cells from "Young" vs "Old" and IPF-affected donors. (A.)** Expression levels of NF-kB activation marker *TNFAIP3* and histone demethylase *JMJD3* mRNA are correlated with each other in AT2 cells after *in vitro* IL1β treatment in mesenchyme-free conditions for 3 days. **(B.)** After 3 days of *in vitro* stimulation with IL1β in mesenchyme-free conditions, there is no significant induction of *KRT5* or *KRT17* mRNA compared to untreated control AT2 cells. **(C-D)** Concomitant treatment with IL1β and NF-kB inhibitor BAY 11-7082 for 3 days shows dose-dependent decrease in both *TNFAIP3* and *JMJD3* mRNA. **(E)** Ingenuity Pathway Analysis (IPA) for differentially expressed genes between NF-kB high and NF-kB low AT2 cells, as determined by their activity scores, in our 4-sample scRNA-seq dataset (n=2 for "Young", n=2 for "Old"). The number of NF-kB high AT2 cells is greater in the "Old" than in the "Young" samples. **(F)** Pseudo-bulk re-analysis of Habermann et al. (Sci. Adv. 2020) scRNA-seq data from IPF Cell Atlas, comparing expression levels between AT2, basaloid (KRT5+/KRT17+) and basaloid (KRT17-/KRT5+) cells as originally annotated by the authors. *JMJD3* expression levels are highest in basaloid cells, whereas there is increase in *KLF5* levels in both basaloid and basal cells. Statistical significance was determined by unpaired t-test for **(B)**.

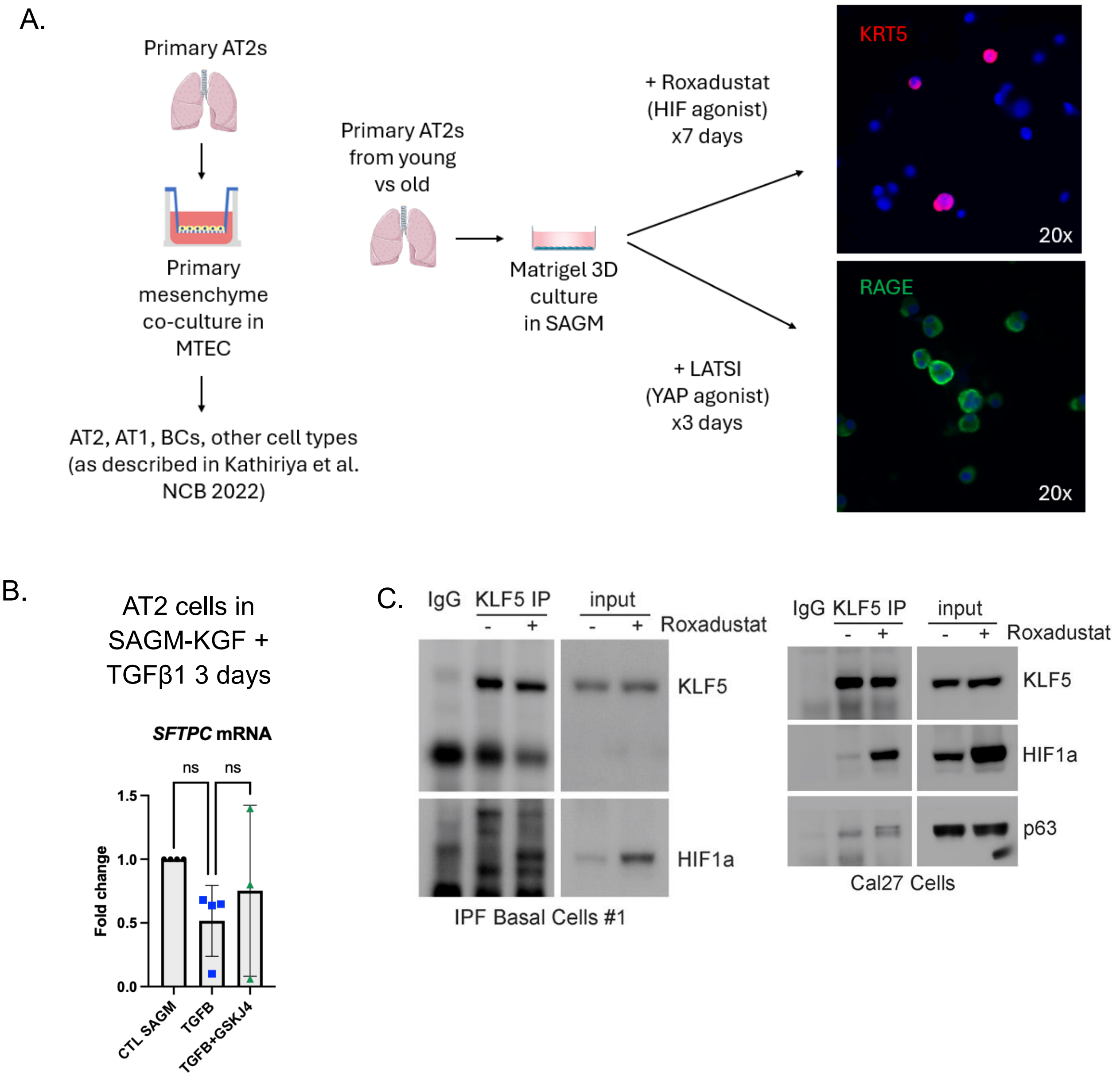


**Supplementary Figure 4**

**Supplementary Figure 4. HIF activation leads to KLF5 re-distribution from AT1-associated loci to basal cell-associated loci in old AT2 cells and antagonizes KRT17 activation by the TGFβ-SMAD2/3 signaling axis, which is specific to a basaloid cell fate. (A)** Representative schematic of how primary human AT2 cells under mesenchyme-free conditions can be forced to acquire either an AT1-like state marked by RAGE expression by treating with a LATS inhibitor, or a basal-like state marked by KRT5 expression by treating with HIF agonist, recapitulating some of the cell types that arise in the fibroblast co-culture organoid system. Importantly, these cells do not appear to be mature AT1 or basal cells at this stage. This system was adapted for KLF5 CUT&Tag (see Figure 4 in the main text). **(B)** Treatment with TGFβ1 +/- GSKJ4 for 7 days does not lead to significant changes in *SFTPC* mRNA expression. **(C)** KLF5 immunoprecipitation in IPF-derived basal cells showing binding with HIF1α after roxadustat treatment. We could not detect p63 protein by Western blot in either the pulldown or cell lysate in these cells. However, we were able to detect p63 protein in a CAL-27 squamous cell carcinoma cell line and observed that p63 co-precipitates with KLF5 with and without HIF1α activation. There may be multiple p63 isoforms of slightly different size bound with KLF5 with HIF1α activation as determined by multiple band appearance. Statistical significance was determined by one-way ANOVA in **(B)**.
